# Midbrain dopamine neurons provide teaching signals for goal-directed navigation

**DOI:** 10.1101/2021.02.17.431585

**Authors:** Karolina Farrell, Armin Lak, Aman B Saleem

## Abstract

In naturalistic environments, animals navigate in order to harvest rewards. Successful goal-directed navigation requires learning to accurately estimate location and select optimal state-dependent actions. Midbrain dopamine neurons are known to be involved in reward value learning^1–13^. They have also been linked to reward location learning, as they play causal roles in place preference^14,15^ and enhance spatial memory^16–21^. Dopamine neurons are therefore ideally placed to provide teaching signals for goal-directed navigation. To test this, we imaged dopamine neural activity as mice learned to navigate in a closed-loop virtual reality corridor and lick to report the reward location. Across learning, phasic dopamine responses developed to visual cues and trial outcome that resembled reward prediction errors and indicated the animal’s estimate of the reward location. We also observed the development of pre-reward ramping activity, the slope of which was modulated by both learning stage and task engagement. The slope of the dopamine ramp was correlated with the accuracy of licks in the next trial, suggesting that the ramp sculpted accurate location-specific action during navigation. Our results indicate that midbrain dopamine neurons, through both their phasic and ramping activity, provide teaching signals for improving goal-directed navigation.

**Highlights:**

- We investigated midbrain dopamine activity in mice learning a goal-directed navigation task in virtual reality
- Phasic dopamine signals reflected prediction errors with respect to subjective estimate of reward location
- A slow ramp in dopamine activity leading up to reward location developed over learning and was enhanced with task engagement
- Positive ramp slopes were followed by improved performance on subsequent trials, suggesting a teaching role during goal-directed navigation

## Results & Discussion

To examine the activity of midbrain dopamine neurons during goal-directed navigation, we designed a task in a virtual reality (VR) corridor. Head-restrained mice were free to self-pace their locomotion on a treadmill, which accordingly updated visual scenes in a closed-loop system^22^ (Figure 1B, Video S1).

**Figure 1.**
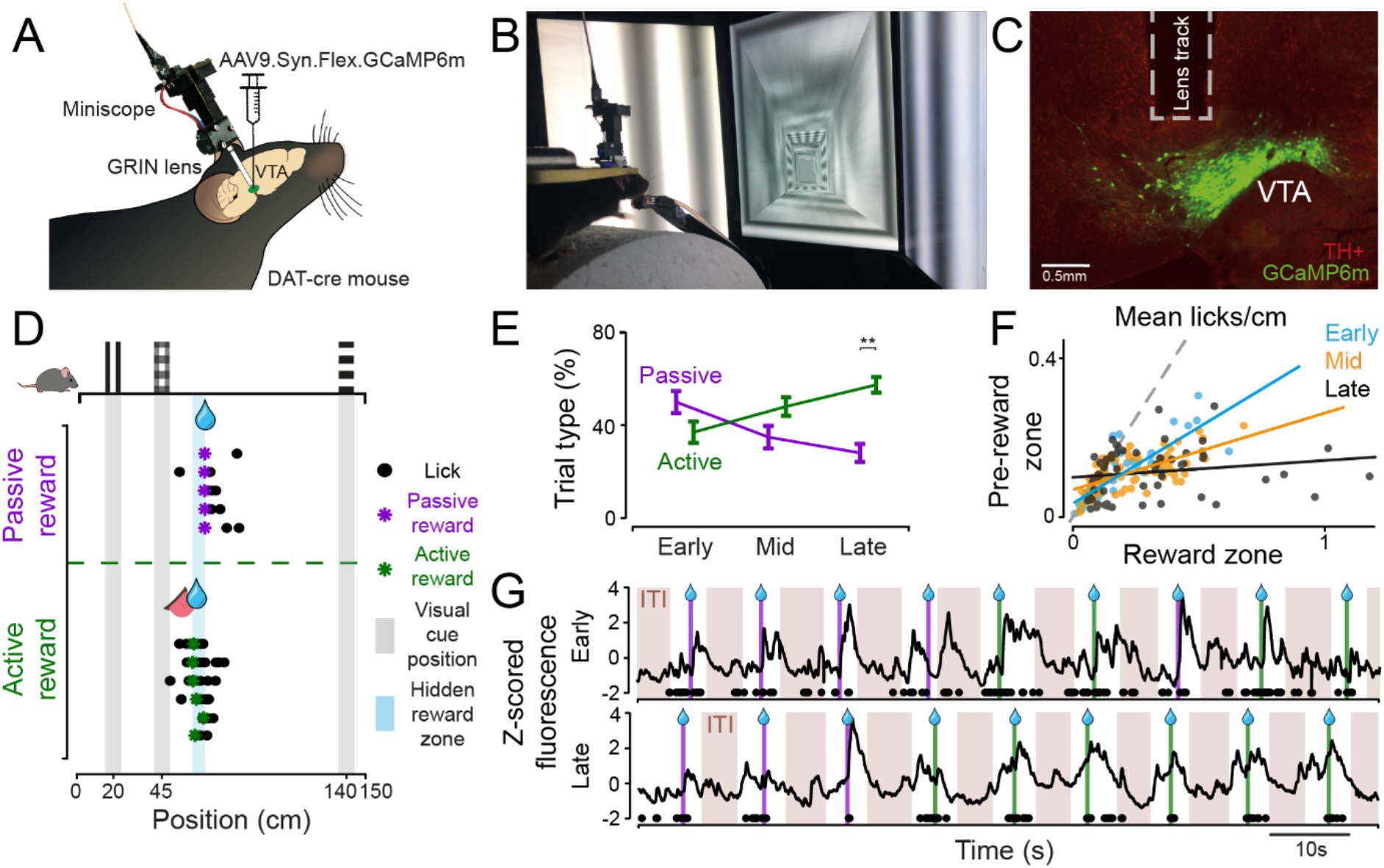
Mice learn to navigate in virtual reality and report the reward location. **A)** DAT-cre mice were injected with AAV9.Syn.Flex.GCaMP6m in the VTA, and implanted with a GRIN lens over the VTA for imaging dopamine activity. **B)** Head-restrained mice performed a navigation task by running on a cylindrical treadmill and virtual corridor displayed on three screens. **C)** Example histological image showing GCaMP6m expression *(green)* in VTA TH+ neurons *(red)* and lens track. **D)** Example behavioural performance shown on a schematic of the corridor with the position of the cues and reward zone, with the licks (*circles*) and reward delivery (*asterisks*) on example trials shown in rows. Licking within reward zone results in active reward delivery (*green*), not licking within reward zone results in passive reward delivery (*purple*). **E)** Mean percentage of passive and active trials across training stages (late-stage, p=0.0078, n=8 animals, Mann-Whitney U test). Error bars indicate standard error. **F)** Comparison of pre-reward lick rate in the reward zone (60-67cm) versus pre-reward zone (50-59cm) at different training stages. **G)** Global fluorescence over several trials from a single animal from early-stage (top) and late-stage (bottom) training stages. Passive (*purple*) and active (*green*) reward deliveries (*lines*), licks (*black circles*), and intertrial intervals (ITIs, *brown shaded regions*) are indicated along the timeline.

A specific region of the virtual corridor had a hidden reward zone, where a lick triggered the delivery of a drop of sweetened water (Figure 1D). The reward zone was not explicitly marked by cues and therefore the mice had to learn to estimate their location based on visual cues presented along the corridor and their own locomotion. If the mice licked within the reward zone, they actively triggered reward delivery (active trial), whereas if mice did not lick in the reward zone, reward was delivered at the end of the reward zone (passive trial). Active trials indicated that the mouse had learned the reward location and reported their subjective estimate of it by licking accurately within the reward zone. We assessed the effects of learning by dividing training sessions into three stages: ‘early’, ‘mid’, and ‘late’ per animal (see Methods). We found that mice performed more active trials and fewer passive trials with increased training (Figure 1E, p=0.0078, Mann-Whitney U test, see Table S1), consistent with their learning the location of the reward zone. Early in training, passive trials indicate that the animal has not yet learned the reward location, whereas later in training, they may indicate task disengagement, erroneous estimation of reward location or attentional lapse. Mice also increased their licking frequency within the reward zone over training (Figure 1F, Figure S2A), indicating that their estimation of the reward location improves over training and they successfully learn to perform the goal-directed navigation task.

As mice learned to perform the goal-directed navigation task in virtual reality, we measured the global activity of midbrain dopamine neurons. We expressed a genetically-encoded calcium indicator (GCaMP6m^23^) using viral transfection in the ventral tegmental area (VTA) of DAT-cre transgenic mice. We implanted a GRIN lens above the VTA, and measured global calcium indicator fluorescence using a Miniscope^24^ (Figure 1, Figure S1). We observed robust phasic responses that followed the reward delivery in individual trials (Figure 1G). Early in learning, dopamine responses mainly appeared after the reward delivery, while later in learning, we observed elevated activity both prior to as well as following the reward delivery, consistent with previous studies^1–13,25^.

Dopamine activity showed sharp, transient increases and decreases following rewarded and unrewarded licks respectively (Figure 2A-B). We calculated the magnitude of the phasic responses as the change in activity from the time of lick to the peak of the response (Figure S3A). Averaged across all sessions, rewarded licks had positive responses (Figure 2A, p<0.0001, Wilcoxon signed rank test), which were larger in active trials compared to passive trials (Figure 2C, p=0.0155, Wilcoxon signed rank test). In contrast, unrewarded licks before reward delivery were followed by a transient suppression in activity (Figure 2B, p<0.0001, Wilcoxon signed rank test). This suppression was followed by a positive phasic response later in the trial when reward was eventually delivered. The magnitude of suppression was similar in both active and passive trials (Figure 2C) but different from responses following rewarded licks (p<0.0001, Mann-Whitney U test). The suppression was consistent with activity suppression we observed in trials where we omitted rewards late in training (Figure S3E-F).

**Figure 2.**
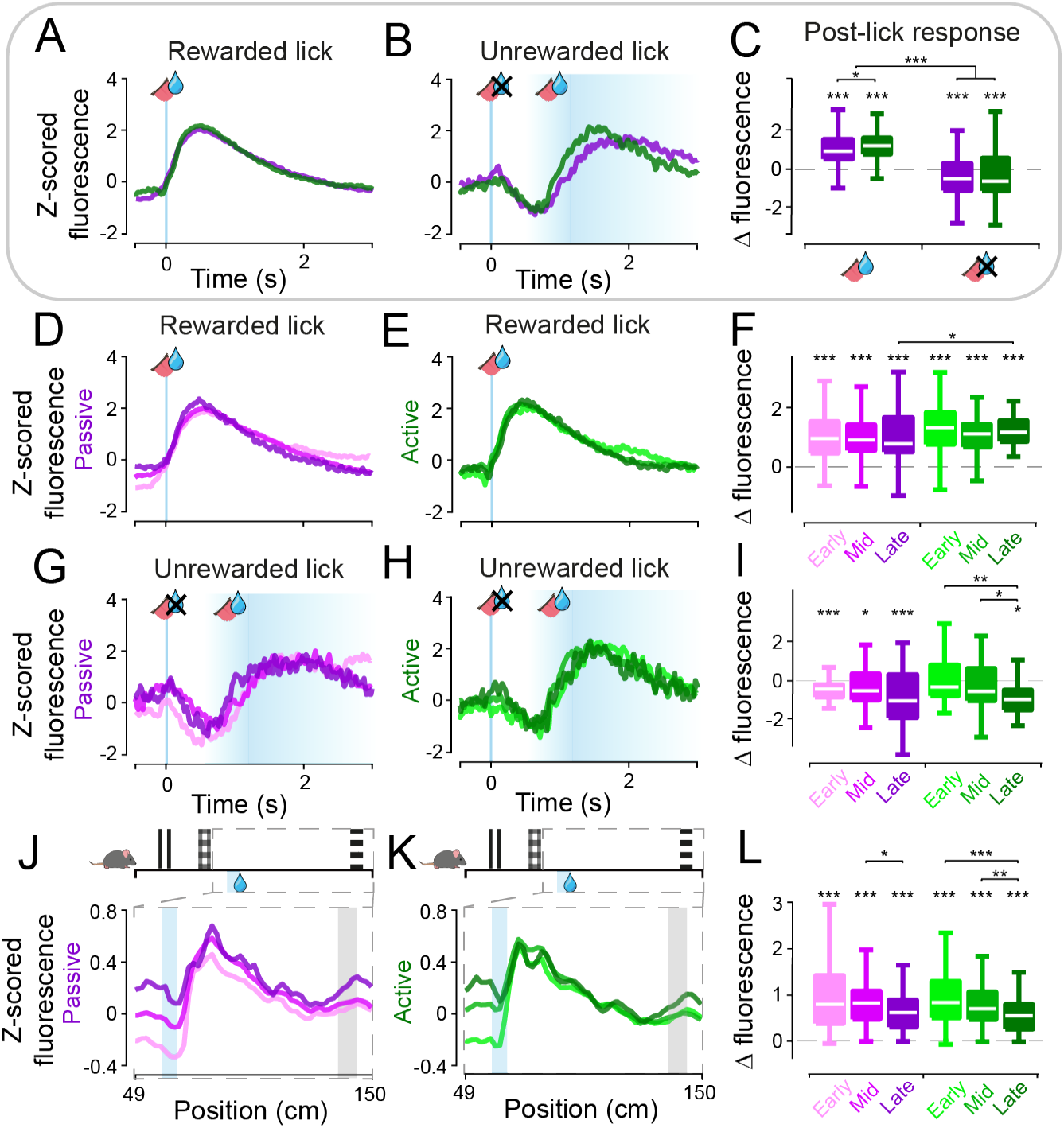
Phasic VTA dopamine activity reflects reward prediction errors. **A-B)** VTA dopaminergic activity as a function of time following rewarded (A) and unrewarded (B) licks for passive (*purple*) and active (*green*) trials. Rewarded licks were taken from trials with no licks prior to reward and the aligned lick is the first lick following reward delivery. Unrewarded licks were taken from trials with one lick >0.5s prior to reward delivery. **C)** Boxplots of change in fluorescence following rewarded (left) and unrewarded (right) licks, measured as maximum difference in the window of 0-0.6s following the lick. Boxplots indicate median across recording sessions (*white*), 25th and 75th percentiles as edges, and whiskers indicate most extreme points (outliers not shown). Asterisks directly above boxplots indicate significant difference from zero (***: p<0.001, **: p<0.01, *: p<0.05; Wilcoxon signed rank test, see Table S1). **D-I)** Same as A-C, split by training stage. **J-L)** Mean dopamine activity as a function of position in the corridor, focused on 49-150cm. Change in fluorescence in L is calculated as the maximum value in the reward window (60-90cm) minus the mean value in the pre-reward window (50-60cm). Change in fluorescence decreases over learning (p<0.05, Mann-Whitney U test, see Table S1).

We also examined how these phasic responses changed over learning (Figure 2D-I). In the time axis, phasic activity following rewarded licks did not change significantly over learning (Figure 2F). For unrewarded licks, we saw the post-lick suppression progressively increase across training in active trials (Figure 2I, e.g. early- vs late-stage: p=0.0078, Mann-Whitney U test, see Table S1), but not in passive trials. Measured along corridor position (Figure 2J-K), we also observed the magnitude of reward responses decrease over training (Figure 2L, e.g. active early- vs late-stage: p<0.001, Mann-Whitney U test).

In summary, the learning-related changes in peri-lick neural activity, particularly when examined in the spatial dimension, are broadly consistent with the reward prediction error term of temporal difference reinforcement learning (RL) models^1,26^. The activity suppression at the time of unrewarded lick further implies that mice in this task have an expectation of reward at the time of lick, reflecting their subjective estimate of the reward location. Finally, differences in phasic responses in active compared to passive trials are reminiscent of previous results showing modulation of dopamine reward prediction errors by factors such as satiety or effort cost^27,28^.

Prior to reward delivery, we observed the development of phasic dopamine activity across training in response to reward-predictive cues, as well as a slow ramp in the pre-reward activity leading up to the reward zone location (Figure 3). Phasic dopamine responses to cues increased over training for both passive and active trials (Figure 3B, e.g. early- vs late-stage: p<0.0001, Mann-Whitney U test). We next examined the slow ramp in pre-reward activity leading up to the reward zone location. Strikingly, we saw that the gradient of the pre-reward ramping activity increased over learning in both passive and active trials (Figure 3C, early- vs late-stage: p<0.0001, Mann-Whitney U test). The gradient of the ramp also appeared to increase earlier in training in active trials compared to passive trials, with the median gradient of the ramp more than doubling for passive trials between mid- and late-stage training (from 0.18 to 0.45 ΔF/cm), while the median gradients were similar for active trials (0.462 and 0.468 ΔF/cm). We saw the same patterns of neural activity emerge over training in trials in which mice did not lick prior to the reward zone, indicating that the pre-reward ramping activity was not caused by licks prior to the reward location (Figure S4). As it has been suggested that dopamine activity could reflect motor vigour^29–33^, we also examined whether locomotor speed could explain the ramping activity. However, ramping dopamine activity did not reflect general locomotor vigour, as mice in our task slowed down on approach to the reward location, while dopamine activity ramped up instead (Figure S2). This resulted in the ramp gradient and change in speed leading up to the reward zone being anti-correlated (Figure S2C).

**Figure 3.**
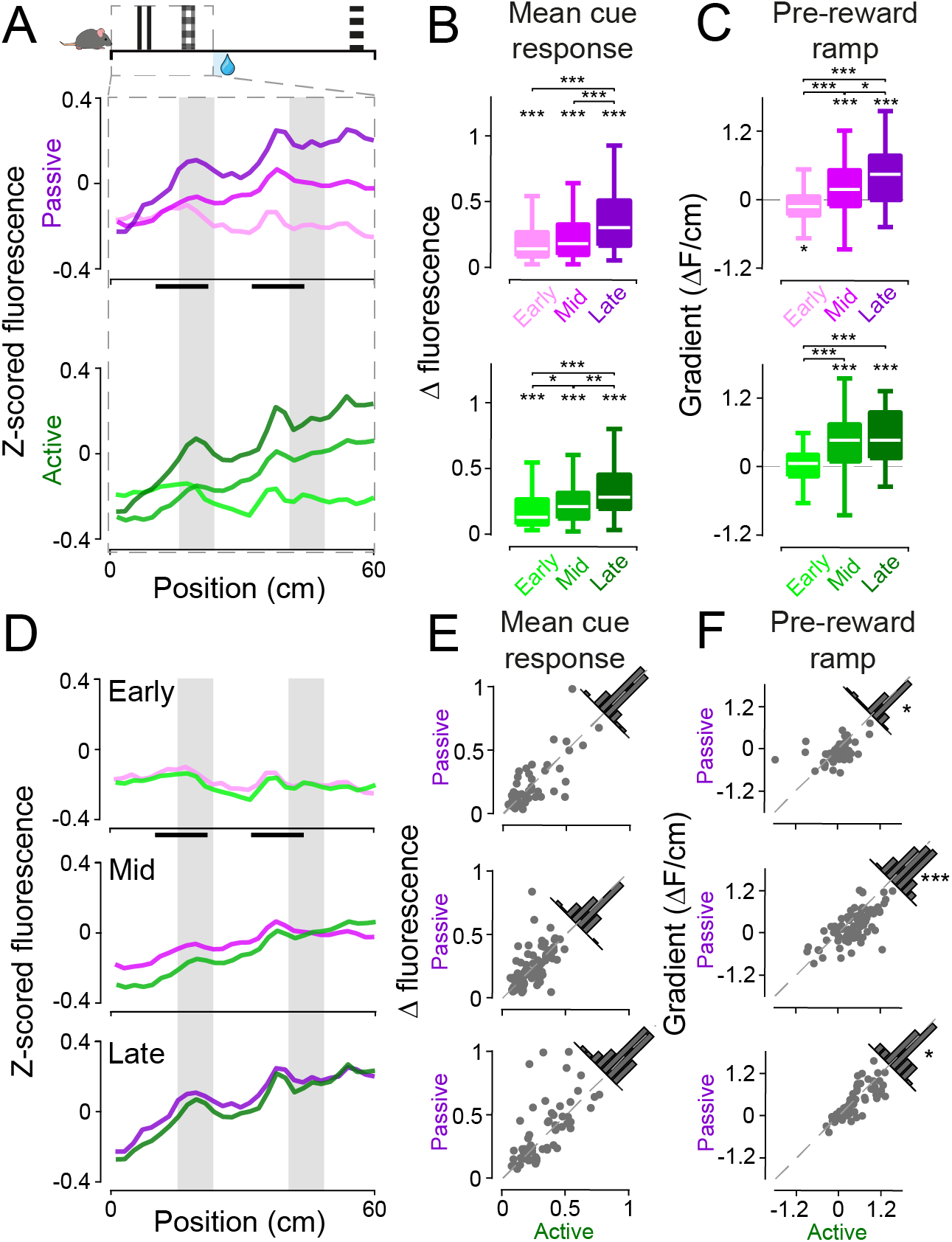
Phasic RPEs and slower pre-reward ramping activity develop over training. **A)** Activity as a function of position in the corridor, focusing on the pre-reward region (0-60cm), split into passive (top) and active (bottom) trials and different training stages. **B)** Boxplots of the mean change in fluorescence in the cue windows indicated by the black bars in A. All distributions are significantly larger than zero (p<0.001, Wilcoxon signed rank test). Change in fluorescence increases over training (p<0.001 for passive, p<0.02 for active, Mann-Whitney U test, see Table S1). **C)** Boxplots of pre-reward ramp gradient, calculated by fitting a line to activity in the 0-60cm window. Asterisks indicate distribution is significantly different from zero (p<0.02, Wilcoxon signed rank test, see Table S1). Pre-reward ramp gradient increases over learning (p<0.05, Mann-Whitney U test, see Table S1). **D-F)** Data shown in A-C, directly comparing passive and active per training stage. Significant differences are found between active and passive ramp gradients at all training stages (Figure 3F, p=0.0243, p<0.0001, p=0.0426 respectively, Wilcoxon signed rank test).

To examine the effect of trial type, we compared cue responses and ramping activity between active and passive trials across the different training stages (Figure 3D). We found that the slope of the pre-reward ramp in active trials was larger than in passive trials at all training stages (Figure 3F, p=0.0243, p<0.0001, p=0.0426 for early-, mid- and late-stage respectively, Wilcoxon signed rank test), while mean cue responses remained similar (Figure 3E). This suggests that the gradient of the ramp was modulated by task engagement.

While ramping of dopamine signals has been observed under certain conditions^4,32,34–41^, its functional role has yet to be agreed upon. Given our observations that the gradient of ramping increased across training, and was higher in active trials (where mice accurately report reward location), we hypothesised that a positive ramp slope might act as a teaching signal to improve performance in reporting the reward zone on a trial-to-trial basis, especially in late-stage training. Individual late-stage trials (with no licking before the reward zone) had pre-reward ramp slopes that ranged from negative to positive (Figure 4A, left). We examined the effect of the sign of pre-reward ramp slope (on trial *n*) on performance in the subsequent trial (trial *n+1*), evaluated by the distribution of licks prior to reward delivery. If a positive pre-reward activity ramp could act as a teaching signal, then we would expect to see increased accuracy of licking in the reward zone in the subsequent trial. Indeed, we found that a positive ramp slope on trial *n* was correlated with more accurate licking in the reward zone on trial *n+1* compared to a negative ramp slope on trial *n* (Figure 4A, p<0.05, Mann-Whitney U test). We also visualised the effect of the positive ramp slope by subtracting the negative slope lick distribution from the positive slope lick distribution (Figure 4B), showing that there is increased licking within the reward zone as a result of the positive slope. The influence on subsequent trials was similar regardless of whether trial *n* was active or passive, albeit with slightly different localisations within the reward zone.

**Figure 4.**
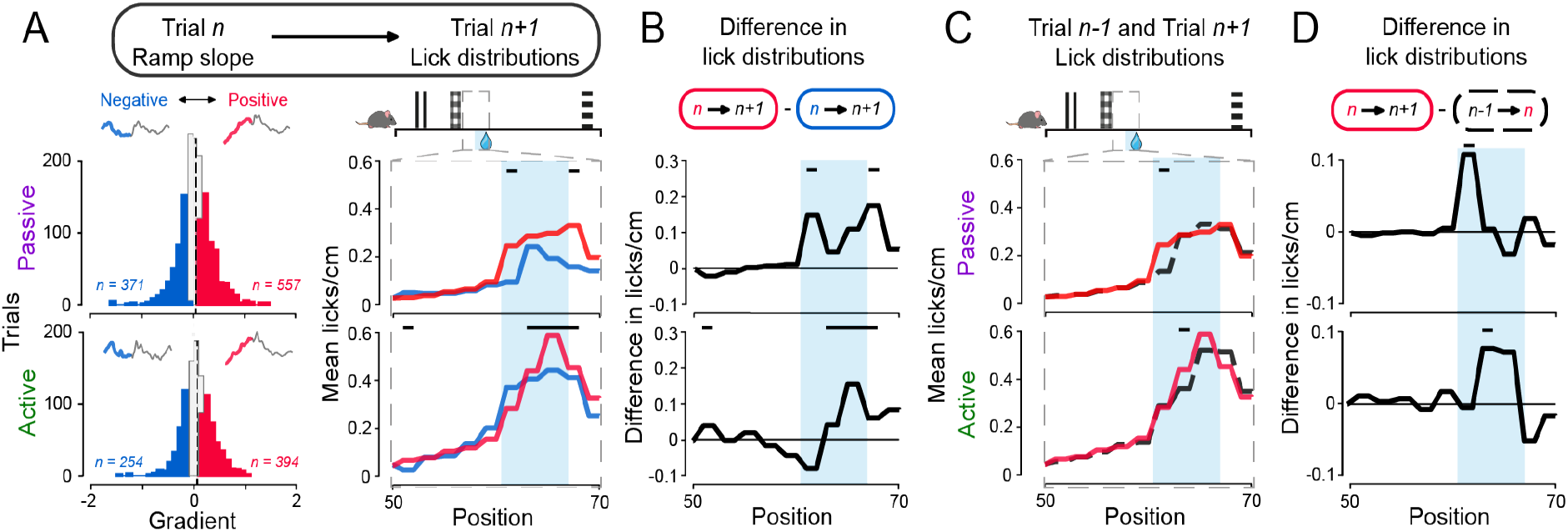
Positive pre-reward ramp slope increases reward zone licking on subsequent trial. **A)** Left: Distributions of pre-reward ramp gradients per late-stage training trial *n* with no licks prior to reward zone, for passive (top) and active (bottom) trials, showing groups of positive (*red*) and negative (*blue*) slope trials. Inset above distributions: mean activity trace for each group of trials. Right: Distributions of pre-reward licks on trials following positive slope trials (*red*) and negative slope trials (*blue*), focusing on 50-70cm in the virtual corridor. Black bars denote significant difference between distributions (p<0.05, Mann-Whitney U test, see Table S1). **B)** Difference between lick distributions shown in A right for passive (top) and active (bottom) trials. **C)** Distributions of pre-reward licks on trials following positive slope trials (*red*) and preceding positive slope trials (*black, dashed*). **D)** Difference between lick distributions shown in C for passive (top) and active (bottom) trials (p<0.05, Mann-Whitney U test, see Table S1).

One possible reason to see this effect of ramp slope on subsequent trials might be that slow fluctuations in behavioural performance cause correlations between neighbouring trials^42^. Such slow fluctuations, however, are not expected to show trial-by-trial improvements in performance. Therefore, to assess trial-by-trial improvements, we used the lick distribution of the trial preceding (trial *n-1*) the positive ramp slope trial *n* as a baseline, and subtracted it from the lick distribution of the subsequent trial *n+1*. If the positive ramp slope improved licking accuracy on a trial-to-trial basis, then we would see an increase in reward zone licking in the lick distribution of trial *n+1* relative to trial *n-1*. As predicted, we saw this effect on subtraction of the trial *n-1* lick distribution from the trial *n+1* lick distribution (Figure 4C-D, p=0.0236 and p=0.0174 for passive and active respectively, Mann-Whitney U test). The effect of the trial *n* positive ramp slope shared similar trial type-specific localisation within the reward zone as we had previously shown (Figure 4B). Interestingly, the same analysis on the effects of reward response sizes on subsequent trials, showed that a small reward response on trial *n* was followed by increased reward zone licking on trial *n+1* when trial *n* was active, but not passive (Figure S5). Together, these data suggest that a positive pre-reward ramp slope acts as a teaching signal, improving the accuracy and frequency of reward location reporting on a trial-to-trial basis.

Overall, our results indicate that both phasic as well as slower ramping of dopamine activity provide teaching signals that can improve the accuracy of goal-directed navigation. We observed the development of positive phasic responses to the reward and the reward-predictive cues, and negative phasic responses following unrewarded licks. In addition, we observed a ramping of the dopamine activity leading up to the reward location, the gradient of which was increased with learning and task engagement. We also saw that a positive ramp slope improved the accuracy of reward location estimation on a trial-to-trial basis.

Our results support the hypothesis that midbrain dopamine neurons provide teaching signals for reward location learning and selecting location-specific actions required to obtain the reward. Phasic dopamine activity represents prediction errors in temporal difference RL models^1,26^, and has been shown to induce learning in various behavioural contexts^5,7,8,11–13^. Phasic dopamine signals incorporate inferred belief state into prediction error computation, reflecting subjective estimates of sensory signals^43^ or reward delivery timing^44^. Our results suggest that phasic dopamine signals also represent belief about the reward location inferred from visual and self-motion cues. Using passive temporal approach tasks in VR, a recent study has shown that ramping dopamine activity mimics RPE^40^. Our results build on this by showing that positive slope ramps improve reward location estimation on a trial-to-trial basis, revealing a role of these ramps in learning in a spatial domain (see Supplementary Discussion). Overall, our results indicate that midbrain dopamine neurons provide teaching signals for goal-directed navigation through both their phasic and slower ramping activity.

## Acknowledgements

We thank Kate Jeffery, Ali Mohebi and Chris Burgess for feedback on the manuscript. This work was supported by an MRC studentship to K.F. (MR/N013867/1), a Sir Henry Dale Fellowship from the Wellcome Trust and Royal Society (213465) to A.L., a Human Science Frontiers Program grant (RGY0076/2018), and a Sir Henry Dale Fellowship from the Wellcome Trust and Royal Society (200501) to A.B.S.

## Author Contributions

Conceptualization, K.F., A.L., and A.B.S.; Methodology, K.F., A.L., and A.B.S.; Investigation, K.F.; Validation, Formal Analysis, Data Curation, K.F.; Writing, K.F., A.L., and A.B.S.; Visualization, K.F., A.L., and A.B.S.; Funding Acquisition, K.F., A.L., and A.B.S.; Resources, A.B.S.; Supervision, A.L., and A.B.S.

## Declaration of Interests

The authors declare no competing interests.

## Methods

KEY RESOURCES TABLE

KEY RESOURCES TABLE
| REAGENT or RESOURCE | SOURCE | IDENTIFIER |
| --- | --- | --- |
| **Antibodies** |  |  |
| Mouse anti-TH | ImmunoStar | Cat # 22941 |
| Alexa Fluor 594 Goat anti-mouse | BioLegend | Cat # 405326 |
| **Bacterial and Virus Strains** |  |  |
| AAV9-Syn-FLEX-GCaMP6m | Addgene | #100838 |
| **Experimental Models: Organisms/Strains** |  |  |
| Mouse: DAT-IRES-cre<br>(Slc6a3<sup>tm1.1(cre)Bkmn</sup>) | The Jackson Laboratory | JAX006660 |
| Mouse: C57BL/6 | Charles River Laboratories | Strain code: 027 |
| **Software and Algorithms** |  |  |
| Bonsai | Lopes et al., 2015<sup>45</sup> | https://bonsai-rx.org/ |
| UCLA Miniscope Bonsai node | Jonathan Newman | https://github.com/jonnew/Bonsai.Miniscope |
| MATLAB 2018a | MathWorks | https://www.mathworks.com/ |
| Custom MATLAB VR code | Saleem et al., 2013<sup>46</sup><br>Saleem et al., 2018<sup>22</sup> | https://github.com/amansaleem/SaleemLab-VR |
| Custom Arduino code for VR interfaces | - | https://github.com/SaleemLab/ArduinoCodes |
| LAS X (Leica Application Suite) confocal microscopy software | Leica Microsystems | https://www.leica-microsystems.com/ |

All procedures were conducted in accordance with the UK Animals Scientific Procedures Act (1986). Experiments were performed at University College London under personal and project licenses released by the Home Office following appropriate ethics review.

### Mouse line creation and maintenance

The DAT-cre transgenic mouse line was started by breeding one male DAT-IRES-cre (Slc6a3^tm1.1(cre)Bkmn^) mice (JAX006660, *The Jackson Laboratory*) with a female C57BL6 mouse (*Charles River*). Following genotypic identification (*Transnetyx*) of DAT-cre offspring, heterozygous DAT-cre breeders were selected and subsequently paired with C57BL6 breeders in order to maintain the colony. Following pregnancy confirmation, males were separated out. Pups were weaned three weeks after birth, earmarked for genotyping, and group-housed in single-sex cages.

All mice were given environmental enrichment, standard chow and water *ad libitum* prior to the experiment. Mice were housed in a colony room at 21.5°C, 45% humidity on a 12h/12h light/dark cycle. Selected experimental mice were single-housed and underwent implant and baseplating surgeries. Following at least 7 days recovery, water-restriction was initiated to increase motivation, with free access to water overnight once every two weeks. Mice were weighed each day, received hydrogel in their home cage following behavioural training to ensure sufficient hydration (>40ml/kg), and had free access to standard chow to maintain their weight between 85-90% of their predicted unrestricted weight. Data from eight mice are presented (5 female, 3 male).

### Miniscope surgeries

Experimental mice underwent two surgeries. In the first surgery, mice were induced with 3% isoflurane and maintained at 1.5%. Lacrilube was applied to eyes to maintain eye moisture, 5% carprofen was administered subcutaneously and the head was shaved. In six animals, 5mg/kg of 2%w/v colvasone (active ingredient: dexamethasone) was administered intramuscularly to reduce inflammation and brain swelling. A craniotomy was performed directly over the VTA of one hemisphere (2 left, 6 right). 600nl of AAV9-Syn-FLEX-GCaMP6m (*Addgene* Plasmid #100838) diluted 1:3 in aCSF was injected at a rate of 50nl/min into the VTA (AP −3mm, ML 0.5mm, DV −4.6mm from dura) and the pipette was left in place for 10 minutes. Following this, for six of the mice, a blunt needle was inserted and lowered between 1.5 and 2mm from dura before being removed. The GRIN lens (*Inscopix* 1050-002179) was then inserted, at an approximate rate of 400-500um/min to a depth around −4.3mm, and secured in place using dental cement (Super-Bond C&B, *Sun Medical*). A custom metal headplate was cemented behind the lens, and a plastic cap (cut-off end of Eppendorf tube) was cemented over the lens for protection. Following recovery, mice were closely monitored and given 20ul Metacam (*Boehringer Ingelheim*) in condensed milk (*Nestlé* Carnation), and high-protein wet food for 3 days post-surgery.

The second surgery was performed 2-3 weeks after the first, to allow for viral expression and inflammation reduction. The mouse was similarly induced, maintained and monitored. Following head-fixation, the protective cap was drilled out and the lens was cleaned. A modified UCLA Miniscope^24^ with an incorporated GRIN lens and with an attached baseplate was lowered to around 100-300um above the implanted lens, and the field of view explored using Bonsai software^45^ and the UCLA Miniscope node (*https://github.com/jonnew/Bonsai.Miniscope*). When the optimal field of view was found, the baseplate was carefully cemented to the skull over the implanted lens. The Miniscope was removed and a protective Delrin cap (*S. Stiteler*, *miniscope.org*) was secured to the baseplate using a set screw.

### Behavioural training and imaging

Mice were handled, water-restricted and acclimatised to head-fixation on a custom Styrofoam wheel^22^ and Miniscope attachment for a few days prior to behavioural testing. Mice were also offered rewards (~1.5-2ul cherry-flavoured Kool-Aid, Kraft Foods), pseudo-randomly delivered, through a lick spout to encourage running and identify putative dopamine reward responses, while monitoring licks using a custom IR sensor. Mice were free to run in the task as much as they desired for about 30 minutes during the dark cycle each day (~5 days/week) on a custom rig, where they were presented with a virtual corridor on three screens (Figure 1B). The three 9.7” screens (LP097QX1-SPAV with 4:3 aspect ratio, controlled by HDMI driver boards) were fixed in portrait mode at 120° from each other, such that they formed half a hexagon, and the mouse was placed at the centre of the hexagon. The mouse’s movements on the wheel were yoked to the visual display using a rotary encoder such that they could only navigate towards the end of the virtual corridor by moving in a forward direction (closed-loop system)^22^ (*github.com/amansaleem/SaleemLab-VR*). The rotary encoder, infrared lick detector, and reward valve (225P011-21, *NResearch*, USA) interfaced with the VR code through an Arduino Leonardo board (*github.com/SaleemLab/ArduinoCodes*). The task used a 150cm long corridor, with a low-contrast white noise pattern along the ceiling, walls and floor (8cm width and height). The visibility of the corridor was limited to 70cm ahead. A full traversal through the corridor is considered a completed trial. Reaching the end (or timing out) initiated an intertrial interval (ITI) where the corridor was replaced with isoluminant grey. The ITI was chosen randomly between 4 and 6s, to ensure that timing between spatial features could not carry past each trial.

As the mice travelled down the corridor, they would pass two distinct patterned cues (8cm wide) on the walls, centred at 20cm and 45cm along the corridor respectively. An unmarked reward zone spanned 60.5cm to 67cm in the corridor. On each trial, a reward was delivered to a spout in front of them. The spout incorporated an infrared sensor to detect licking. The exact location of the reward zone was not indicated by any cue and instead had to be estimated by the mouse based on prior cues and actions. If the mouse did not lick in the reward zone, then it would passively receive the reward at the end of the zone (passive trials). However, if it licked within the zone, then reward delivery was actively triggered (active trials), and therefore delivered earlier than in the passive trials (Figure 1D). The mouse could then continue down the virtual corridor and pass a final, non-reward-predictive patterned cue (centred at 140cm) before the end of the corridor was reached (grey screen). If the mouse did not reach the end of the corridor within 30 seconds, the trial was terminated (timed out). A pre-reward licking threshold was also imposed to reduce licking and indicate the mouse’s estimation of the reward location. This was gradually reduced over training (following the mouse’s natural inhibition of excessive licking in incorrect locations) to approximately 8-10 licks in late-stage training. If the mouse exceeded this threshold prior to reward delivery, the trial was terminated.

For comparison of data across training, training sessions were split into three stages: early, mid and late training. Early and late stages were defined as the first and last quartile of sessions respectively for each animal, with the rest being classified as mid-training.

Calcium fluorescence was detected using a Miniscope through an implanted GRIN lens (Figure 1A) which acted as a proxy for dopamine neuron activity. A custom Bonsai workflow^45^ and Miniscope node (*https://github.com/jonnew/Bonsai.Miniscope*) were used to acquire images (at 15Hz) and calculate global calcium signal. Mice had a mean of 24 training sessions over the course of the experiment. In early to mid-training, active trials were incentivised by offering slightly larger rewards (~2-3ul) until the mice demonstrated the ability to repeatedly perform active trials (as judged by the experimenter), at which point the active trial reward volume was decreased to be the same as the passive trial reward volume. Later in training, 6 of the 8 mice had reward omission trials introduced pseudo-randomly in 5-7% of the trials in each session, where no reward was delivered but the mouse still traversed the corridor.

### Data pre-processing

Imaging data collected using Bonsai was imported into Matlab for pre-processing. Rarely, unstable signal was produced by Miniscope movement or power surges or lapses. Two types of signals were considered unstable. The first type was fluorescence that exceeded or fell below a threshold of 1.5 standard deviations away from the mean fluorescence across the whole training session. The second type was when fluorescence surrounding the first type (±100ms) exceeded half of the difference between the maximal fluorescence and mean fluorescence, therefore constituting an ‘upswing’ or ‘downswing’ of a large transient. Following removal of these two types of unstable signal, a photobleaching curve was fitted across the entire session using Matlab *polyfit* (2^nd^ order) and subtracted from the fluorescence trace. The trace was then corrected for jitter by subtracting the lower 10% quantile baseline using a 60s window. The resulting signal was then aligned to the virtual corridor times. Trials with unstable signals were removed from subsequent analysis.

To be included in further analysis, trials, sessions and animals had to fulfil certain criteria. Aborted trials (time-outs, too many licks before reward or experimenter-terminated), and trials with unstable signal were excluded from analysis. Sessions were included if they had >50% trials with at least one lick, and >10% active trials. Four animals were excluded from further training as they did not show visible reward responses to random reward, and were later confirmed to have mistargeted GRIN lens placement. One animal was excluded due to a visual defect (cataract), and another one was excluded as it did not learn the task (based on having <50% of the sessions containing active trials). Subsequent analysis was performed on data that met these conditions (eight animals). Fluorescence was z-scored prior to comparisons across sessions or animals.

### Data analysis

Behaviour during training was assessed through binned mean licks/cm across the virtual corridor, as well as through the speed of the wheel rotation. Phasic responses to the cues were calculated as the maximum minus the minimum values within each 12cm cue window (10-22cm, 32-44cm and 132-144cm respectively). Phasic responses to the reward were calculated as the maximum value in the reward window (60-90cm) minus the mean of the activity within the pre-reward window (50-60cm). Ramp gradient was calculated as the gradient of a fitted linear line (Matlab *polyfit*, 1^st^ order) to the fluorescence in the window 0-60cm.

Rewarded lick traces (Figure 2) included only trials that did not have any licks before the reward zone, to avoid contamination of the signal by prior licks. Rewarded lick traces were also averaged across each animal before averaging over all animals to counter the appearance of an electrical artifact that was presented in two animals when reward was delivered during active trials. Unrewarded lick traces included only trials that had only one lick prior to the reward zone that was at least 0.5s before reward delivery. For comparison with reward omission trials, the data in Figure S3 only includes sessions that contained omission trials. Suppression gradient was calculated as the mean of the gradients of fitted lines (Matlab *polyfit*, 1^st^ order) between the fluorescence at zero and the minimum fluorescence in the second half of the window of fluorescence being examined (here −2.5 to 3s around the lick, so minimum value between 0.25 and 3s) for each trial. Paired data was tested for differences using the Mann-Whitney U test (Matlab *ranksum*), as were tests of difference from zero, while the Wilcoxon signed rank test (Matlab *signrank*) was used to test for differences between different training stages.

Ramp slopes were classified as positive or negative by considering all ramp slopes across all trials, sessions and animals and taking the most positive third as ‘positive’ and the most negative third as ‘negative’. For analysis of the effect of ramp slope on trial *n* on lick distribution on trial *n+1*, only trials with no licks before the reward zone were considered to eliminate any contamination of signal by licks. Lick distributions in Figure 4 were calculated as licks/cm smoothed from 2cm bins. Figure 4C-D analysis used lick distribution from trial *n-1* for normalisation to isolate trial-by-trial effect^42^.

### Histology

Mice were deeply anaesthetised using 3.5% isoflurane, injected with a lethal dose of pentobarbital (Euthatal, *Boehringer Ingelheim*) intraperitoneally and transcardially perfused with 1X phosphate-buffered saline (PBS) followed by 10% formalin solution. Following perfusion, the brain was extracted and placed in 10% formalin for short-term storage. Prior to sectioning, the brain was placed into a 30% sucrose solution until it sank, for cryoprotection. The brain was then mounted upright in OCT (*Sakura* FineTek) and 40um slices were made using a cryostat (*Leica* CM1850 UV). Slices were washed five times in 1X PBS before overnight incubation on a rotating platform at room temperature in primary solution: 1:5000 mouse anti-TH (*ImmunoStar* 22941) in PBS-T (0.4% Triton in 1X PBS), to label tyrosine hydroxylase-positive (including dopamine) cells. The following day, slices were washed five time in 1X PBS before a 2-hour secondary incubation, in 1:1000 Alexa Fluor 594-goat anti-mouse (*Biolegend*) in PBS-T. Slices were then washed five times in 1X PBS before being mounted and allowed to dry. Mounting medium with DAPI (Vectashield, *Vector Laboratories*) was added to stain cell bodies, before adding the coverslip and sealing with nail polish. Slices were then imaged at 10x magnification (Figure S1) using a confocal microscope (*Leica* DMi8) and LAS X software (*Leica*).

## Supplemental Information

### Supplementary Discussion

#### Midbrain dopamine neurons and spatial navigation

Our study revealed the presence of reward prediction error signals during goal-directed navigation in the dopaminergic neurons of the VTA. Previous work using manipulations of midbrain dopamine neural activity established their role in spatial learning, such as in place preference assays^14,15^ and in spatial memory tasks via their effects on hippocampal plasticity, place fields and ensemble reactivation^16–21^. However, their well-known function in signalling reward prediction error (RPE), which was established in the temporal domain^1,26,48^, has not yet been precisely demonstrated in spatial tasks. This is partly because most studies of midbrain dopamine neurons during navigation have used freely-moving animals in tasks that don’t have precise temporal features or behavioural readouts. By performing experiments in virtual reality (VR), we were able to create a navigation task with high temporal precision and a precise readout of the animal’s estimate of reward location through licking^22,49^. This allowed us to measure neural responses to precise events such as cues, rewards, and licks, and establish the presence of reward prediction errors during spatial learning.

Our implementation of VR also has the advantage of being closed-loop. If progression through the virtual corridor was simply presented as a video of movement at a predefined speed, irrespective of the animal’s own movements, spatial components could not be differentiated from an equivalent temporal task^40^. In contrast, our closed-loop task gives control of movement (and corresponding visual scenes) to the animal, simulating more naturalistic navigation. We were therefore able to characterise neural activity as a function of spatial position, which revealed ramping activity of VTA dopaminergic activity along the corridor leading up to the reward location, similar to dopamine ramps observed in animals navigating real environments^37,38^.

Another key advantage of our task is the ability to assess the engagement of animals in the task. We were able to split trials into ‘active’ and ‘passive’, based on whether the animal actively triggered reward delivery by licking in the reward zone or not. Once animals had learned how to perform the task, this allowed us to define two degrees of task engagement, and revealed that both the phasic post-rewarded lick responses and the ramping of pre-reward dopaminergic activity were greater when the animal was engaged and actively reporting the reward location.

#### Function of ramping dopaminergic activity

Pre-reward dopamine ramping has been observed^4,32,34–41,50^ in tasks that tend to have two particular features: first, reward is distant, often requiring several seconds to be attained, and second, progress towards the reward can be tracked, such as through sensory cues or action sequences^4,32,34–41,50^. These criteria are both naturally met during goal-directed navigation. The pre-reward ramp has been reported in both VTA dopamine neuron activity^34,39–41,52^ and dopamine release into downstream striatum^4,32,35–38^ and across species. We also observed pre-reward ramping in VTA dopaminergic neural activity in our goal-directed navigation task in mice. This ramping activity developed over learning, was modulated by task engagement and was correlated with improvements in reward location estimation on a trial-to-trial basis. However, the functional significance of the pre-reward ramping is currently up for debate.

One hypothesis suggested for pre-reward ramping activity is that it represents reward prediction error (RPE)^40^. Our results are largely consistent with this hypothesis. Specifically, we observed the pre-reward ramp develop over learning, increasing in gradient while phasic RPE signals to reward-predictive cues increased in magnitude. We also found that positive slope ramps are followed by increased reward zone licking on the subsequent trial on a trial-to-trial basis, indicating that a positive slope ramp is correlated with improved reward location estimation and performance, much like RPE improves performance on a trial-to-trial basis^11,42^.

An alternate hypothesis is that dopamine ramps reflect value based on the animal’s inferred current state^4,32,51^. Many of our results are consistent with this hypothesis. In addition, during navigation, the value of the animal’s current state may not be distinguishable from estimated goal proximity^37,41,52^. These representations reflect ongoing behaviour, however, our finding that dopamine ramps are followed by improved reward location estimation on a trial-by-trial basis suggests that ramping has a teaching role for improving future behaviour. However, these functions may not be mutually exclusive, as ramping could be present in multiple neuronal populations or the ramp could represent a multiplexed signal^39,50^.

General motivation and vigour are other functions that have been suggested to be fulfilled by ramping dopamine^32,37,38,51^, such that ramping dopamine might serve to motivate the animal to locomote and act vigorously as it approaches the goal and harvests the reward. However, our data are inconsistent with this hypothesis, as our animals slow down on approach to the reward zone in our task, while VTA dopamine neuron activity ramps up (Figure S2). This presumably reflects a speed-accuracy trade-off required to accurately lick in the reward zone while also moving rapidly, maximising the number of trials (and therefore amount of reward harvested) during each training session. This is in contrast to many other studies of dopamine function, where instead the animal aims to perform the task as quickly as possible, and therefore self-paced movement speed correlates with VTA dopaminergic activity^37,53–55^. Our task is therefore able to delineate vigorous action from accurate action, and indicates that ramping dopamine activity does not reflect behavioural vigour or a general motivation to exert more effort (or energy) in a scalar fashion (i.e. run faster). An alternate explanation is that accurate reward zone licking might require more cognitive effort. Slowing down is therefore a strategy to increase accuracy, and ramping dopamine activity might represent the specific goal-directed motivation to slow down and/or lick accurately. This highlights the problem of ambiguity in how motivation and effort are interpreted in different studies, and the need for replication with respect to task requirements, motor actions performed, and the subpopulations and anatomical localisation of the dopamine neurons recorded.

Dopamine neural activity has also been suggested to encode action initiation and aspects of movement^3,33,39,50,53–56^. However, we find no relation between ramping VTA dopamine neuron activity and action (lick) initiation, as ramps are also seen in trials where no licking occurs prior to the reward zone (Figure S4). We also did not find transients preceding or peaking at lick time, but only phasic responses following the lick which show RPE based on whether the lick was rewarded or not (Figure 2). In terms of locomotion, we found that our mice mostly ran continuously, mediating their speed along the virtual corridor. As they slowed down on approach to movement, speed and ramping activity were negatively correlated (Figure S2). Our results indicate that VTA dopamine neuron activity does not reflect action initiation or speed in our goal-directed navigation task.

Overall, our data show a ramping of VTA dopamine neuron activity that is consistent with encoding of RPE, state value and goal proximity. It could also be construed as supporting a specific, goal-directed motivation signal, but not a general behavioural activation, movement or vigour signal. Our analysis of trial-by-trial improvement in reward location estimation following ramping activity favours a role in learning, similar to RPE encoding. Together, we conclude that VTA dopamine neurons provide teaching signals, through both their phasic and slower ramping activity, for goal-directed navigation.

#### Supplementary Figures

**Video S1. Video of mouse behaviour.** Video showing experimental setup and the behaviour of a mouse over two trials of the goal-directed navigation task in virtual reality. The mouse was head-fixed over a cylindrical treadmill with a Miniscope attached to its head. The mouse ran at a self-defined pace through a virtual corridor displayed on three screens, and licks in the lick port to receive reward. Reward delivery is heard as a click as the reward valve opens. The video is annotated to visually show when the reward is delivered (*blue dot*). After the corridor has been traversed, the screens display iso-luminant grey for an intertrial interval (4-6s), after which the animal is transported back to the start of the corridor to initiate the next trial.

**Figure S1.**
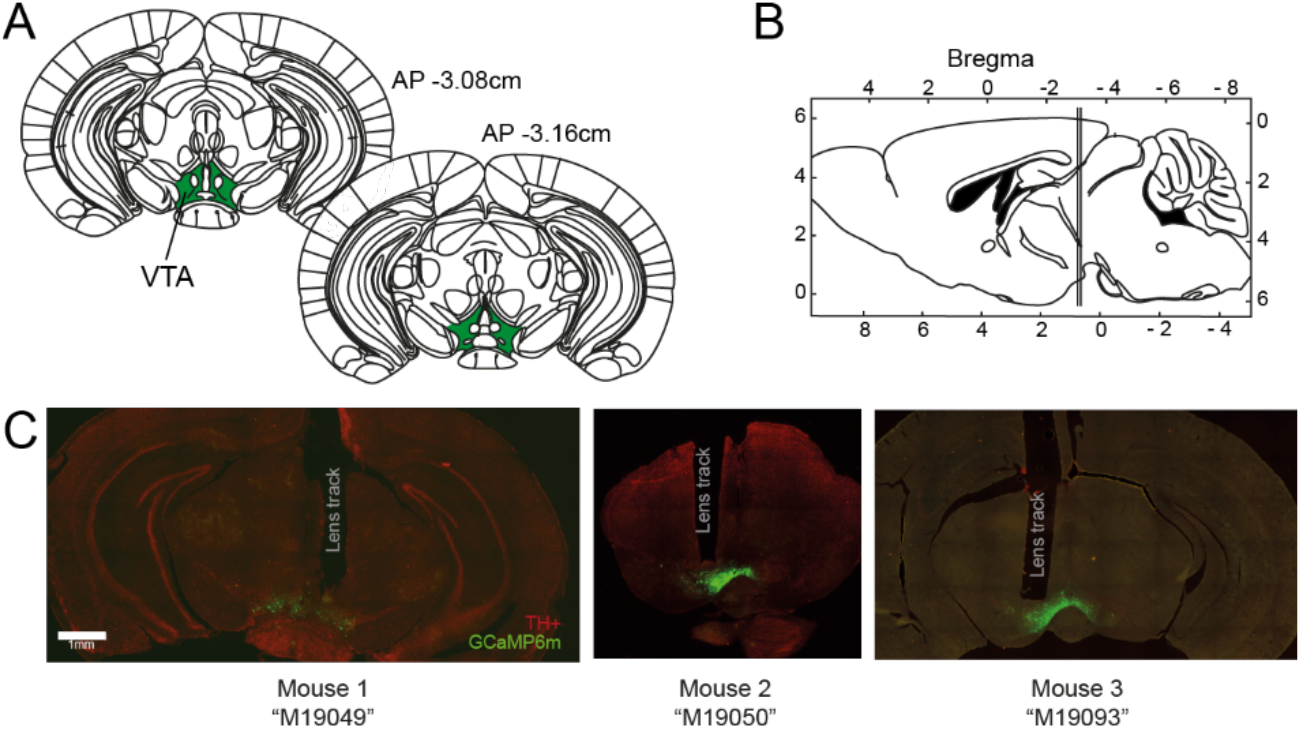
Histology from example mice. **A)** Figures 56 and 57 from Paxinos and Franklin (2001)^47^, showing diagram of horizontal section of mouse brain at −3.08cm at −3.16cm from bregma, with VTA highlighted in green. **B)** Inset of Figures 56 and 57 from Paxinos and Franklin (2001)^47^ showing diagram of sagittal section of mouse brain, with sections at −3.08cm and −3.16cm from bregma indicated. **C)** Example histology from three mice, showing GCaMP6m (*green*) and tyrosine hydroxylase (TH) staining (*red*).

**Figure S2.**
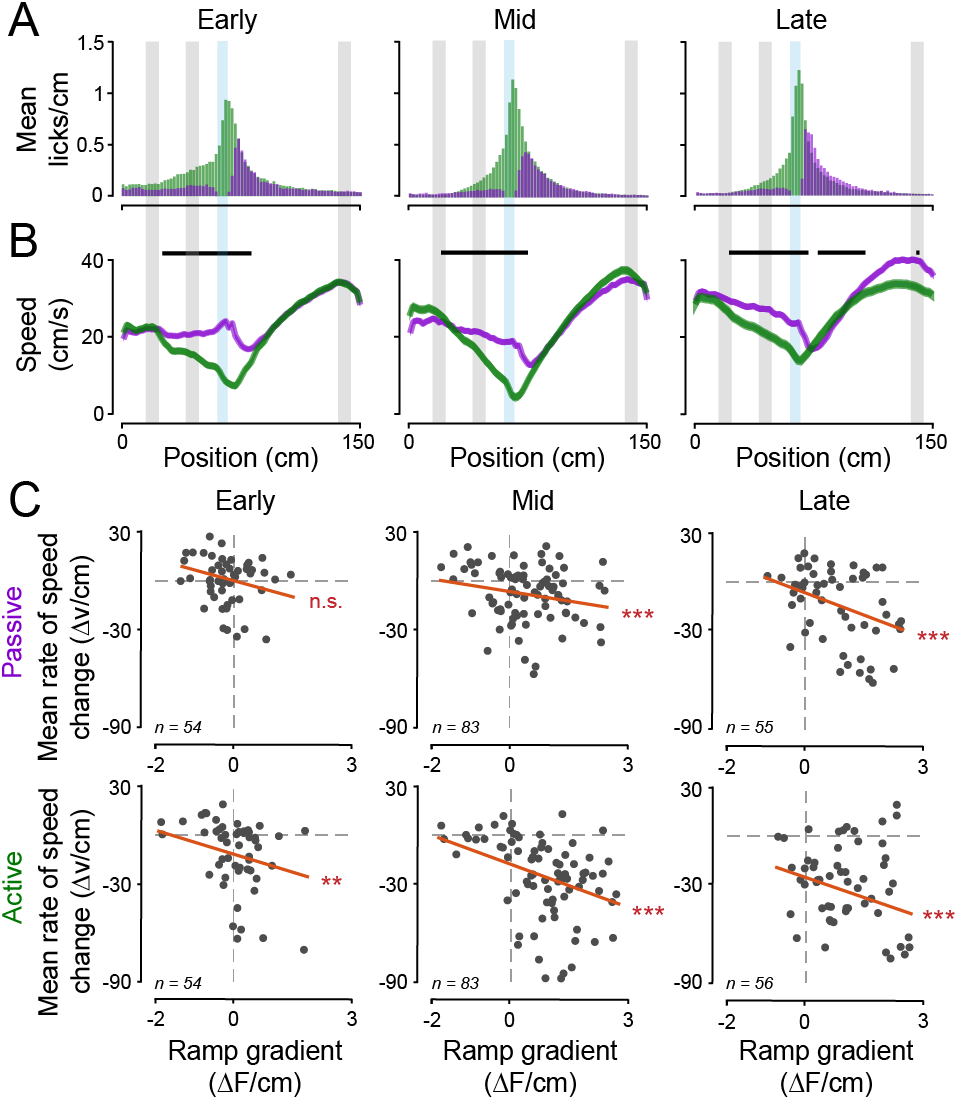
Behaviour and speed analysis. **A)** Mean lick distribution per training stage for active (*green*) and passive (*purple*) trials. **B)** Mean speed profile per training stage for active and passive trials. Black bars indicate significant difference between active and passive speed profiles for positions indicated (early-stage: n=55, p<0.001 for bins 30, 34, 38-78cm, p<0.01 for bins 26-28, 32, 36, 80cm, p<0.05 for bin 82cm, mid-stage: n=83, p<0.001 for bins 28-74cm, p<0.01 for bin 26cm, p<0.05 for bins 10, 22, 24, 76cm, late-stage: n=55, p<0.001 for bins 34-70, 80, 84, 90-92cm, p<0.01 for bins 24-32, 78, 82, 86-88, 94-98, 104cm, p<0.05 for bins 22, 72, 100-102, 108, 140-142cm, Wilcoxon signed rank test). **C)** Gradient of fitted line to change in speed over pre-reward distance (mean rate of pre-reward speed change, calculated as change in speed per cm) plotted against gradient of fitted line to pre-reward calcium activity per session for each training stage for passive (top) and active (bottom) trials. Line fitted to points using Matlab *polyfit* shown in red. Wilcoxon signed rank test for passive early (p=0.2134, n=54), mid (p<0.0001, n=83), late (p<0.001, n=56) and active early (p=0.0063, n=54), mid (p<0.0001, n=83) and late (p<0.0001, n=55).

**Figure S3.**
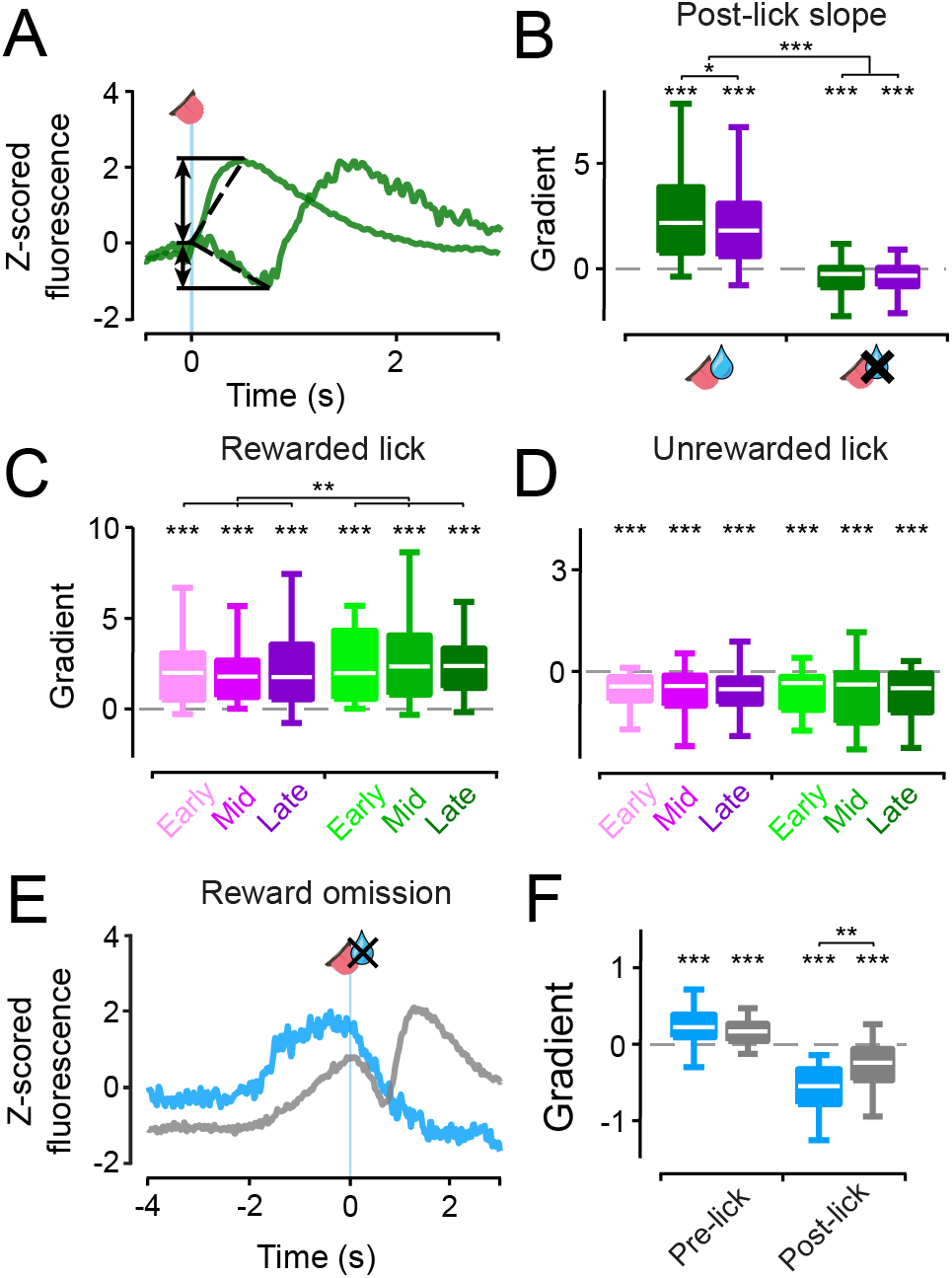
Gradient of post-lick responses following rewarded and unrewarded licks, and reward omission. **A)** Schematic of how change in fluorescence and lines are fitted to mean of activity traces from active trials following rewarded and unrewarded licks (as shown in Figure 2A-B). **B)** Boxplots of gradient of slope following lick for both rewarded and unrewarded licks, as shown in Figure 2A-B. Asterisks directly above boxplots indicate significant difference from zero (p<0.0001 for all, n=188 (passive, rewarded), n=176 (active, rewarded), n=187 (passive, unrewarded), n=193 (active, unrewarded), Wilcoxon signed rank test). Difference between gradient of slope for rewarded licks in active and passive trials is significant (p=0.0376, n=176, Mann-Whitney U test), as is the gradient of the slope following rewarded licks vs unrewarded licks (p<0.0001, n=191, Mann-Whitney U test). **C-D)** Boxplots of gradient of post-lick response following rewarded and unrewarded licks, split by training stage, corresponding to trace in Figure 2D-E. All boxplots are significantly different from zero (p<0.0001 for all, n=54, 82, 52 (passive, rewarded, early-mid-late respectively), n=45, 78, 53 (active, rewarded, early-mid-late respectively), n=48, 57, 36 (passive, unrewarded, early-mid-late respectively), n=31, 43, 33 (active, unrewarded, early-mid-late respectively), Wilcoxon signed rank test). The mean gradient of the post-rewarded lick response is significantly greater in active compared to passive trial (p=0.0075, n=176, Wilcoxon signed rank test). **E)** Mean activity traces averaged over sessions from reward omission trials aligned to time of lick in reward zone (*blue*) and unrewarded licks with only one lick occurring before the reward zone (and reward following later in the trial) from the same sessions as the omission trials (*grey*). **F)** Comparison of pre-lick gradient (calculated by fitting a line to activity in the window of −3s to time of aligned lick) and post-lick gradient for the traces shown in E. Pre-lick gradients are significantly greater than zero, whereas post-lick response gradients are significantly lower than zero (p<0.0001 for all, n=37 (omission, pre-lick), n=50 (non-omission, pre-lick), n=35 (omission, post-lick), n=39 (non-omission, post-lick), Wilcoxon signed rank test). The post-lick gradients are also significantly different between the reward omission trials and the unrewarded trials (p=0.0427, n=35, Wilcoxon signed rank test).

**Figure S4.**
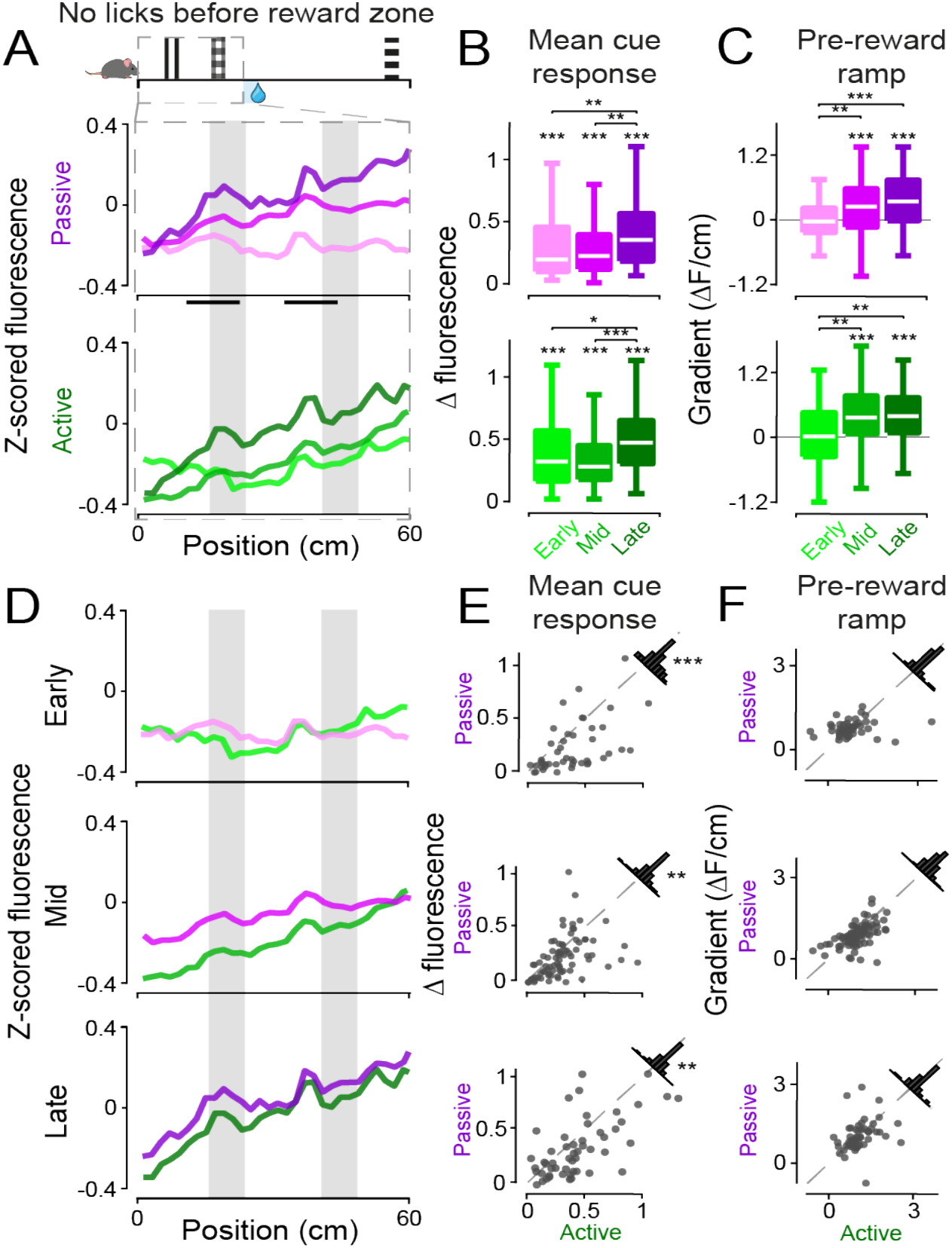
Pre-reward activity in trials that had no licking prior to the reward zone. **A)** Mean activity traces from trials where no licking occurred prior to the reward zone, focusing on the pre-reward corridor region from 0-60cm, split into different training stages. Black bars indicate position windows where cue responses are calculated for use in B. **B)** As in Figure 3B, boxplots of the mean of maximal change in fluorescence for the two cue windows indicated by the black bars in A, split by training stage. Values from each trial are averaged over each session. All distributions are significantly larger than zero (p<0.0001 for all, n=54, 83, 54 (passive, early-mid-late respectively), n=45, 78, 54 (active, early-mid-late respectively), Wilcoxon signed rank test). Change in fluorescence for passive trials is significantly different between early- and late-stage training, as well as mid- and late-stage training (p=0.003 and p=0.005 respectively, n=54, Mann-Whitney U test). For active trials, change in fluorescence is significantly different between early- and late-stage training, as well as mid- and late-stage training (p<0.0335 and p<0.0001 respectively, n=45, 54, Mann-Whitney U test). **C)** Boxplots of the mean pre-reward ramp gradient, calculated by fitting a line to activity in the 0-60cm window. Asterisks above mid- and late-stage boxplots for active and passive trials indicate distribution is significantly greater than zero (p<0.001 for all, n=83 (passive, mid), n=54 (passive, late), n=78 (active, mid), n=54 (active, late), Wilcoxon signed rank test). Pre-reward ramp gradient for passive trials is significantly different between early- and mid-stage training, and early- and late-stage training (p=0.0049 and p<0.001 respectively, n=54, Mann-Whitney U test). For active trials, ramp gradient is significantly different between early- and mid-stage training as well as early- and late-stage training (p=0.0013 and p=0.0014 respectively, n=45, Mann-Whitney U test). **D-F)** Same data shown in A-C, but directly comparing passive and active for each training stage. For E, significant differences are found between active and passive mean change in fluorescence per session for the two cues at early-, mid- and late-stage training (p<0.001 and n=45, p=0.0042 and n=58, p=0.0026 and n=73 respectively, Wilcoxon signed rank test).

**Figure S5.**
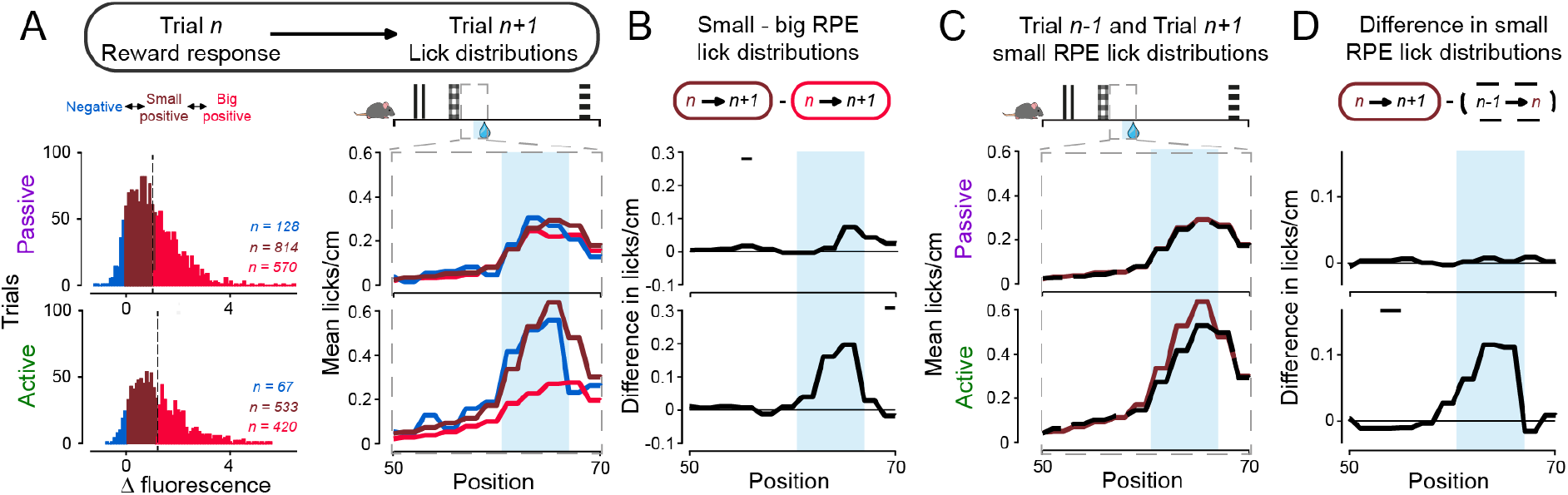
Trial-to-trial analysis of reward response. **A) Left:** Distributions of post-reward delivery reward responses (RPEs) per late-stage training trial *n* with no licks prior to reward zone, for passive (top) and active (bottom). Groups are big positive RPE trials (*red*), small positive RPE trials (*brown*) and negative RPE trials (*blue*). **Right:** Distributions of pre-reward licks on trials following big positive RPE trials (*red*), small positive RPE trials (*brown*) and negative RPE trials (*blue*), focusing on 50-70cm in the virtual corridor. **B)** Difference between lick distributions shown in A) right for passive (top) and active (bottom) trials. Black bars denote a significant difference between distributions (for passive p=0.0253, n=570 for 55-56cm, for active p=0.0229, n=420 for 69-70cm, Mann-Whitney U test). **C)** Distributions of pre-reward licks on trials following small positive RPE trials (*brown*) and preceding small positive RPE trials (*black, dashed*). **D)** Difference between lick distributions shown in C for passive (top) and active (bottom) trials (for active p=0.0048, n=531 for 63-64cm, Mann-Whitney U test).

**Table S1:** List of statistical tests shown in figures. [**M-W:** Mann-Whitney U test (paired); **W(u):** Wilcoxon signed rank test (unpaired); **W(p):** Wilcoxon signed rank test (paired)]

| Fig | Variables | p-value | n | Test |
| --- | --- | --- | --- | --- |
| 1E | Late-stage percentage passive trials, late-stage percentage active trials | 0.0078 | 8 | M-W |
| 2C | Passive post-rewarded lick responses | 5.9094e-23 | 188 | W(u) |
| 2C | Active post-rewarded lick responses | 8.8950e-24 | 176 | W(u) |
| 2C | Passive post-unrewarded lick responses | 1.4454e-07 | 141 | W(u) |
| 2C | Active post-unrewarded lick responses | 2.2765e-10 | 107 | W(u) |
| 2C | Passive post-rewarded lick responses, active post-rewarded lick responses | 0.0155 | 194 | W(p) |
| 2C | Post-rewarded lick responses, post-unrewarded lick responses | 2.7494e-22 | 194 | W(p) |
| 2F | Early-stage passive post-rewarded lick responses | 6.3378e-09 | 54 | W(u) |
| 2F | Mid-stage passive post-rewarded lick responses | 7.2065e-12 | 82 | W(u) |
| 2F | Late-stage passive post-rewarded lick responses | 1.1845e-05 | 52 | W(u) |
| 2F | Early-stage active post-rewarded lick responses | 6.2459e-07 | 45 | W(u) |
| 2F | Mid-stage active post-rewarded lick responses | 1.5766e-10 | 78 | W(u) |
| 2F | Late-stage active post-rewarded lick responses | 3.8267e-09 | 53 | W(u) |
| 2F | Late-stage passive post-rewarded lick responses, late-stage active post-rewarded lick responses | 0.0406 | 52 | W(p) |
| 2I | Early-stage passive post-unrewarded lick responses | 2.8253e-04 | 48 | W(u) |
| 2I | Mid-stage passive post-unrewarded lick responses | 0.0012 | 57 | W(u) |
| 2I | Late-stage passive post-unrewarded lick responses | 6.5155e-04 | 36 | W(u) |
| 2I | Late-stage active post-unrewarded lick responses | 0.0044 | 33 | W(u) |
| 2I | Early-stage active post-unrewarded lick responses, late-stage active post-unrewarded lick responses | 0.0078 | 31, 33 | M-W |
| 2I | Mid-stage active post-unrewarded lick responses, late-stage active post-unrewarded lick responses | 0.0352 | 43, 33 | M-W |
| 2L | Early-stage passive post-reward responses | 2.2765e-10 | 54 | W(u) |
| 2L | Mid-stage passive post-reward responses | 2.7928e-15 | 83 | W(u) |
| 2L | Late-stage passive post-reward responses | 2.9637e-10 | 55 | W(u) |
| 2L | Early-stage active post-reward responses | 1.9244e-10 | 54 | W(u) |
| 2L | Mid-stage active post-reward responses | 3.3498e-15 | 83 | W(u) |
| 2L | Late-stage active post-reward responses | 7.7267e-10 | 56 | W(u) |
| 2L | Mid-stage passive post-reward responses, late-stage passive post-reward responses | 0.0237 | 83, 55 | M-W |
| 2L | Early-stage active post-reward responses, late-stage active post-reward responses | 4.0522e-04 | 54, 56 | M-W |
| 2L | Mid-stage active post-reward responses, late-stage active post-reward responses | 0.0039 | 83, 56 | M-W |
| 3B | Early-stage passive mean cue responses | 1.6257e-10 | 54 | W(u) |
| 3B | Mid-stage passive mean cue responses | 2.5034e-15 | 83 | W(u) |
| 3B | Late-stage passive mean cue responses | 1.1076e-10 | 55 | W(u) |
| 3B | Early-stage passive mean cue responses, late-stage passive mean cue responses | 1.0094e-05 | 54, 55 | M-W |
| 3B | Mid-stage passive mean cue responses, late-stage passive mean cue responses | 1.0300e-04 | 83, 55 | M-W |
| 3B | Early-stage active mean cue responses | 1.6257e-10 | 54 | W(u) |
| 3B | Mid-stage active mean cue responses | 2.5034e-15 | 83 | W(u) |
| 3B | Late-stage active mean cue responses | 7.5475e-11 | 56 | W(u) |
| 3B | Early-stage active mean cue responses, mid-stage active mean cue responses | 0.0169 | 54, 83 | M-W |
| 3B | Early-stage active mean cue responses, late-stage active mean cue responses | 5.3701e-05 | 54, 56 | M-W |
| 3B | Mid-stage active mean cue responses, late-stage active mean cue responses | 0.0015 | 83, 56 | M-W |
| 3C | Early-stage passive ramp gradients | 0.0185 | 54 | W(u) |
| 3C | Mid-stage passive ramp gradients | 1.2954e-04 | 83 | W(u) |
| 3C | Late-stage passive ramp gradients | 3.9987e-09 | 55 | W(u) |
| 3C | Early-stage passive ramp gradients, mid-stage passive ramp gradients | 1.4195e-05 | 54, 83 | M-W |
| 3C | Early-stage passive ramp gradients, late-stage passive ramp gradients | 6.0579e-09 | 54, 55 | M-W |
| 3C | Mid-stage passive ramp gradients, late-stage passive ramp gradients | 0.0212 | 83, 55 | M-W |
| 3C | Mid-stage active ramp gradients | 2.9572e-09 | 83 | W(u) |
| 3C | Late-stage active ramp gradients | 5.4663e-09 | 56 | W(u) |
| 3C | Early-stage active ramp gradients, mid-stage active ramp gradients | 8.6079e-08 | 54, 83 | M-W |
| 3C | Early-stage active ramp gradients, late-stage active ramp gradients | 2.7530e-08 | 54, 56 | M-W |
| 3F | Early-stage active ramp gradients, early-stage passive ramp gradients | 0.0243 | 54 | W(p) |
| 3F | Mid-stage active ramp gradients, mid-stage passive ramp gradients | 1.3368e-05 | 83 | W(p) |
| 3F | Late-stage active ramp gradients, late-stage passive ramp gradients | 0.0426 | 55 | W(p) |
| 4A/B | Late-stage zero licks before reward passive positive slope subsequent trial lick distribution, late-stage zero licks before reward passive negative slope subsequent trial lick distribution: bin 61-62cm, bin 67-68cm | 0.0116, 0.0009 | 568, 392 | M-W |
| 4A/B | Late-stage zero licks before reward active positive slope subsequent trial lick distribution, late-stage zero licks before reward active negative slope subsequent trial lick distribution: bin 63-64cm, bin 65-66cm, bin 67-68cm | 0.0283,<br>0.0032,<br>0.0403 | 398,<br>262 | M-W |
| 4C/D | Late-stage zero licks before reward passive positive slope subsequent trial lick distribution, late-stage zero licks before reward passive positive slope previous trial lick distribution: bin 61-62cm | 0.0236 | 568,<br>564 | M-W |
| 4C/D | Late-stage zero licks before reward active positive slope subsequent trial lick distribution, late-stage zero licks before reward active positive slope previous trial lick distribution: bin 63-64cm | 0.0174 | 398,<br>398 | M-W |

## Notes

### Competing Interest Statement

The authors have declared no competing interest.

